# A marked enhancement of a BLOC-1 gene, *pallidin*, associated with somnolent mouse models deficient in histamine transmission

**DOI:** 10.1101/2022.05.04.490425

**Authors:** Laurent Seugnet, Christelle Anaclet, Magali Perier, Jean-François Ghersi-Egea, Jian-Sheng Lin

## Abstract

Histamine (HA) and orexin (Ox, or hypocretin) neurons act distinctly and synergistically in wake control. A double knock out mouse genotype lacking both HA and Ox shows all sleep disorders of human narcolepsy. We identified in this mouse brain a sharp upregulation of a BLOC-1 gene, *pallidin* that is associated with dramatic changes in the balance of cholinergic and aminergic systems in mice and an enhanced sleep in drosophila. This study demonstrates potential sleep disorders-associated compensatory mechanisms with *pallid* as a novel biomarker.

---

The maintenance of wakefulness requires a complex brain arousal network made up of diverse neurotransmitters and neuropeptides. Using knock out (KO) mouse models we have previously shown that histamine (HA) and orexin (Ox, also called hypocretin) neurons act distinctly and synergistically in terms of wake control^1,2^. An impaired histaminergic neurotransmission is associated with sleepiness in animal models and human sleep disorders while the lack of Ox neuropeptides constitutes a direct cause of narcolepsy, a neurological disease characterized by sleepiness and cataplexy^1,2^. We have generated a double KO mouse genotype lacking both HA and Ox (*Histidine decarboxylase-Orexin* KO, referred to as *HO*-KO) which shows all phenotypes of human narcolepsy such as sleepiness, hypersomnia, sleep-onset rapid eye movement and cataplexy-like episodes, EEG hypersynchronization and marked obesity^3^. This mouse strain constitutes therefore a complete murine model of narcolepsy. In order to identify the consequences on gene expression of sleep disorders and to uncover novel molecular and cellular processes involved in sleep-wake control, we performed transcriptomic profiling in the frontal cortex of *HO*-KO mice and their wild type littermates.

We identified differentially expressed genes in this double mutant mouse that potentially reflect unidentified mechanisms controlling sleep and wakefulness (Supplemental table 1). Figure 1A shows a subset of these genes confirmed by independent quantitative PCR (QPCR). The non-protein coding Hdc transcript was highly upregulated likely because of a negative feed-back regulation between HA release and *Hdc* expression^4^ (Figure 1B). We obtained similar findings for the *Ox* non-protein coding transcript (Figure 1B, middle). Interestingly this regulation was not uniform in the brain, indicating local compensatory regulatory mechanisms on neurotransmission. The expression of genes not previously associated with sleep-wake regulation were also affected, notably *pallidin* (also called *BLOC1S6*), a gene coding a major subunit of the Biogenesis of Lysosome-related Organelles Complex-1 (BLOC-1), that displayed a massively enhanced expression (> 900%), in the frontal cortex as well as in the hypothalamus and thalamus (Figure 1B). To determine whether the upregulation of *pallidin* results from a HA or Ox deficiency or both, we performed QPCR in single KO mice lacking either *Hdc* or *Ox*. We found that enhanced *pallidin* expression occurred only in *Hdc*-KO mice and not in *Ox*-KO mice, indicating a HA-dependent upregulation. We further found that *pallidin* expression was also highly upregulated in KO mice lacking postsynaptic HA H1-receptor, confirming that the lack of HA neurotransmission plays a key role (Figure 1C) in the regulation of this gene. The Ox transmission does not seem to affect *pallidin* expression which was unchanged in Ox KO mice. Yet, *pallidin* expression was markedly higher in the *HO*-KO than in *Hdc*-KO mice (Figure 1C), suggesting that the abnormal sleep phenotypes associated with Ox deficiency could contribute to the *pallidin* upregulation.

**Figure 1.**
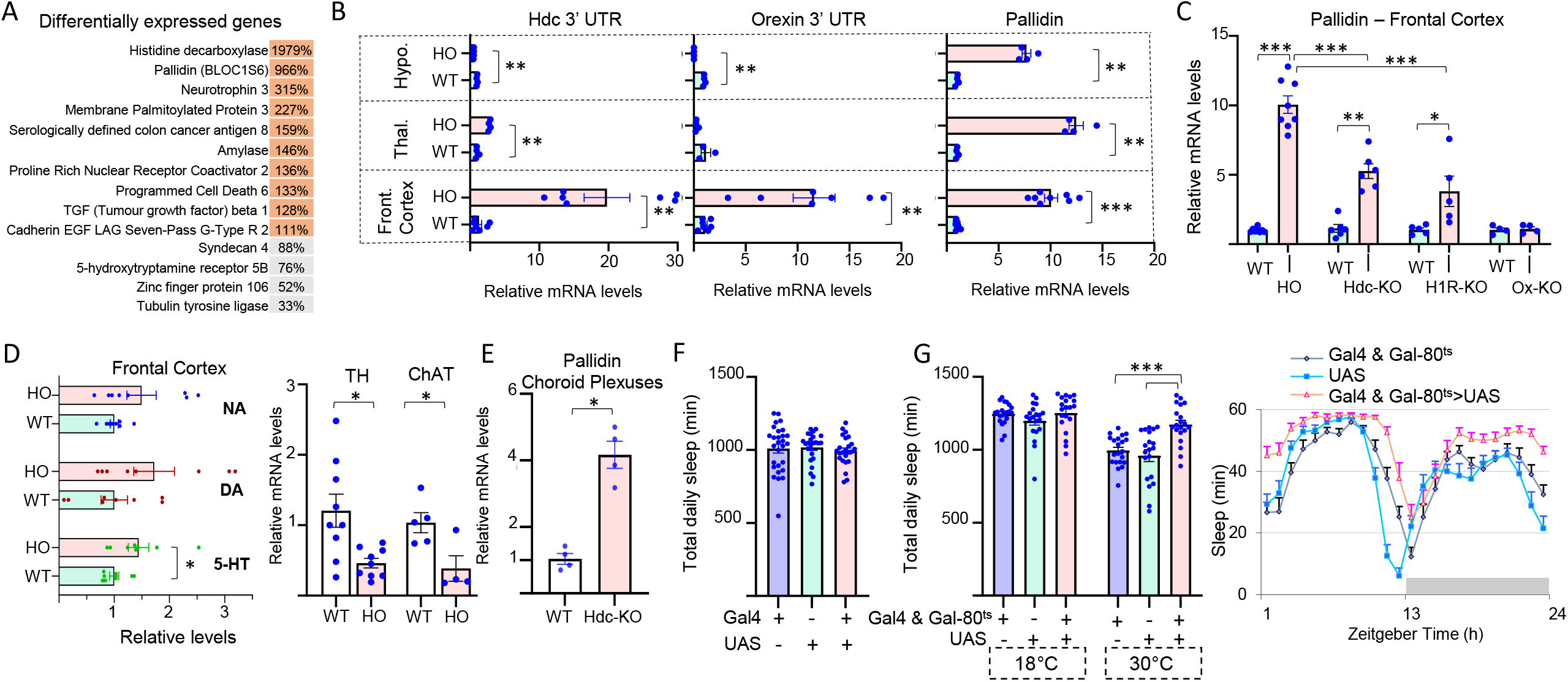
A, a selection of differentially expressed genes in *Histidine decarboxylase-Orexin* KO *i*.*e*., *hdc-Ox or HO*-KO mice, confirmed by QPCR. Relative expression as % of controls is shown (N=3-7, p<0,05). B, Relative expression levels as % of controls in the hypothalamus (top row), thalamus (middle row) and frontal cortex (bottom row) of *HO*-KO mice, for the 3’ untranslated part of the truncated transcripts of *Hdc* and *Ox* in the KO. Right: expression of *pallidin* in the same brain regions. N=4-8. C, Relative expression levels of *pallidin* transcripts in the frontal cortex of HO-KO compared to *hdc, Histamine H1 Receptor* (*H1R)* and (*Ox*) single KO mice. N=4-8. D, left: levels of Noradrenaline (NA), Dopamine (DA) and Serotonin (5-HT) as measured by HPLC in the frontal cortex of *HO*-KO mice, right: relative transcripts levels for the *tyrosine hydroxylase* (TH) and *Choline acetyltransferase* (ChAT) genes in the frontal cortex of *HO*-KO mice (N=4-9). E, Relative expression levels of *pallidin* transcripts in the choroid plexuses of *hdc* single KO mice. F-G, Overexpression of *pallidin* in *Drosophila*. The experimental flies are in pink, the genetic controls are in blue. N=20-21. F, Overexpression of *pallidin* in neurons, the elav-GeneSwitch construct was used as Gal4. G, Overexpression of *pallidin* in all glial cells using the Gal4-UAS system. The Repo-Gal4 transgene was combined with the Tub-Gal80^ts^ to inhibit Gal4 activity at 18°C. Flies were transferred at 30°C to induced the expression of *pallidin*. Left: total daily sleep, right: hourly sleep curve of the first 24h spent at 30°C. *: p<0,05; **: p<0,005; ***: p<0,0005. See supplemental methods for additional details and statistics.

The BLOC-1 complex is involved in protein trafficking among different endosomal compartments and has been linked to schizophrenia and cognitive performance^5^. Indeed, BLOC-1 genes play important roles in neuronal functions and in particular neurotransmission^5^. In addition, it has been reported that *pallidin* could regulate transport at the blood-brain-barrier and thus impact monoamine synthesis, and in particular that of serotonin^6^. In line with this hypothesis, we found a marked increase in serotonin levels (Figure 1D), a change in expression of the *5HT-5B* receptor gene (Figure 1A), and a significant decrease in mRNA levels for *Tyrosine Hydroxylase* in the frontal cortex of *HO*-KO mice (Figure 1D). *Choline acetyltransferase* mRNA levels are also decreased in *HO*-KO mice indicating that the cholinergic system is affected (Figure 1D). HDC activity and HA transmission are present in non-neuronal cell types of the brain, notably in the different blood-brain interfaces, including the choroid plexuses which form the blood-CSF barrier, where HA affects gene expression^7,8^. We thus evaluated *pallidin* mRNA expression in the choroid plexuses of *Hdc*-KO mice, and found that it was upregulated in these non-neuronal cells (Figure 1E) as in the frontal cortex (Figure 1B).

A study based on locomotor activity suggested that *pallidin* could be involved in sleep-wake regulation in the mouse^9^, yet, direct evidence is lacking. The role of *pallidin* in sleep-wake regulation remains, therefore, to be explored, in particular using conditional, cell type specific KO approaches. Yet, such attempt is currently extremely challenging in mammals given the broad expression of *pallidin* in numerous cells types in the brain and periphery and currently the lack of flexible tools to selectively vary *pallidin* expression. We thus turned to the *Drosophila* model, which is intensively used in sleep-wake research. All the genes involved in BLOC-1 function are conserved in *Drosophila*, with similar interactions and functions as those identified in mammals^10^. We tested neuronal and non-neuronal function for *pallidin* using pharmacological or heat inducible transgenes Gal4-UAS systems to overexpress *pallidin* (Figure 1F-G). In *Drosophila*, glial cells regulate neurotransmission and control the exchanges between the brain and circulating fluid, the hemolymph, fulfilling a function similar to the blood-brain-barrier in mammals. We found that overexpression of *pallidin* in neurons did not produce any detectable effect (Figure 1F) while that in the glia using a heat inducible system significantly enhanced sleep (Figure 1G).

*Pallidin* appeared among the most dramatically up-regulated genes in the brain of mice deficient for HA transmission and characterized by a somnolent phenotype. Moreover, we found that over-expression of *pallidin* in *Drosophila* glia results in increased sleep time. Therefore, *pallidin* upregulation in *HO*-KO and *Hdc*-KO is likely linked to the altered control of sleep-wake in these somnolent mice. Gene expression dosage appears to be a critical factor in the stability of the BLOC-1 complex^5^ both in mammals and *Drosophila*. If such is the case in the mammalian blood-brain interfaces, the upregulation of *pallidin* may significantly modulate the functionality of BLOC-1. The signaling pathway leading to this transcriptional response together with its potential impact on sleep-wake regulation remain to be identified. HA transmission has been shown to regulate the blood brain interfaces^7^ and ependymal cells, the circumventricular organs and the choroid plexuses are in close proximity with HA terminals. Interestingly, all of the latter cells display dramatically enhanced *c-fos* expression in response to acute pain induced by formalin injection, and in some cases even to saline injection, in *hdc* KO mice^8^, suggesting that the lack of HA transmission enhances transcriptional responses. In *Drosophila*, HA neurons modulate sleep-wake via chloride-gated channels, and no metabotropic HA receptor has so far been identified, indicating that the interactions between the HA system and *pallidin* differ from those in mammals. Nevertheless, we found that increasing *pallidin* expression in *Drosophila* glial cells increased sleep, an effect potentially linked to the transport of monoamines precursors, since monoamines such as serotonin and dopamine are major evolutionary conserved sleep-wake regulators and since in particular serotonin levels and *TH* mRNA were markedly affected in HO KO mice.

The differentially expressed genes identified in this study suggest widespread compensatory mechanisms upon impaired HA transmission. The unexpectedly sharp upregulation of *pallidin* might constitute part of such adaptive mechanisms allowing the brain to compensate for the lack of HA. Alternatively, this upregulation could be directly linked to the sleepiness caused by the deficient HA transmission. These questions open future investigations. In any case, *pallidin*, easily quantifiable by PCR, might constitute a biomarker of sleepiness in animals and in humans. Similarly, studies on whether and how the identified changes in gene expression are involved in the regulation of neurotransmitter systems, notably acetylcholine, monoamines, and/or metabolism, should open new avenues in sleep-wake research.

## Acknowledgments

This work was supported by Integrative Physiology of the Brain Arousal Systems (CNRL, Inserm U1028 -CNRS UMR5292 – UCBL1), ANRs NarConX and Histawake. We gratefully thank Sandrine Parrot (NeuroDialyTics, Inserm U1028, UCBL1) and Nathalie Turle-Lorenzo (LNC, UMR 7291, Aix Marseille Université) for performing the HPLC experiments. We thank Hideo Akaoka for skilled dissections of the mouse brains.

