## Supplemental information for "A marked enhancement of a BLOC-1 gene, *pallidin*, associated with somnolent mouse models deficient in histamine transmission"

**Supplementary Materials and methods**

**Transcriptomics and quantitative real time PCR**

Transcriptomics. Previously described KO mice for Hdc[1] and Ox[2,3] in C57BL/6 genetic background were crossed to generate the HO double KO mice. 3 *HO* KO mice and 3 wild type littermates were sacrified at 4-5 months old. The frontal cortex was dissected and used as a starting material. CodeLink whole mouse genome arrays were used to assess gene expression. Whole genome profiling and analysis were carried out by the ProfileXpert facility.

Quantitative real time PCR . Total RNA from mouse frontal cortex or from micro-dissected choroid plexuses was extracted using the RNeasy Microkit with DNase treatment (Qiagen). Reverse transcription was then performed using 500 ng of total RNA, with Bio-Rad Laboratories' iScript cDNA Synthesis Kit. Quantitative real-time PCR was performed on a 7900HT Fast Real-Time PCR System using FAST SYBR Green Master Mix (AppliedBiosystems). The levels of the reference gene glyceraldehyde-3-phosphate dehydrogenase (Gapdh) transcript were used to normalize the potential amount variation of sample cDNAs added to each reaction. The relative expression ratio was then determined using the 2ΔΔCt calculation method. Mean ± SEM is plotted, a *t*-test was used for statistical comparison between the WT and KO conditions. ANOVA followed and posthoc Tuckey tests were used to compare *HO*- *Hdc*- and *H1R*- KO in Figure 1 C.

Sequences of the primers: *Ox* (exon 2, still present ion the KO[2]) forward: tctacgaactgttgcacgga, reverse: ccatttaccaagagactgaca; *Hdc* (exon 12, still present in the KO[1]) forward: atgcaagagtgcctgtgctt, reverse: agcatgcccgcttaaacttc; *Pallidin* forward: gccactggcagggtttcccaca, reverse: gccactggacggtctatcccacca; *Tyrosine Hydroxylase* forward: aagggcctctatgctaccca, reverse: gccagtccgttccttcaaga, *Choline acetyltransferase* forward: gcctcatctctggtgtgctt, reverse: atacagagaggctgccctga, *GAPDH* forward: actgagcaagagaggccctta, reverse: tatgggggtctgggatggaa.

**Analysis of monoamine content using HPLC**

The cortical tissues from OH-KO mice were very quickly microdissected and weighted, then kept at -80°C before monoamine extraction. On the day of the extraction, they were slightly thawed and suspended in 10 µL per mg of tissue of ice-cold 0.1 mol/L perchloric acid containing 1.34 mmol/L EDTA and 0.05%, w/v sodium bisulfite and sonicated for 2 × 15 s, then the homogenates were centrifuged at 16,000 × g for 20 min at +4°C. Monoamines (dopamine, serotonin and noradrenaline; as in [4], except an injection volume reduced to 0.1 µL) were quantified using high performance liquid chromatography coupled to electrochemical detection. Results were normalized to the mean of the

WT condition. Mean ± SEM is plotted, Mann-Whitney tests were used for statistical comparison between the WT and KO conditions.

***Drosophila* experiments**

Fly husbandry. Flies were raised on standard food containing inactivated yeast, cornmeal, agar, molasses, sucrose and maintained at 25°C, 60% humidity in a 12hr: 12hr Light: Dark (LD) cycle. The following stocks obtained from the Bloomington Drosophila Stock Center : *Canton S*, repo-Gal4, Tub-Gal80$^{ts}$, PBac{WH}Pldn$^{f05716}$. The PBac{WH}Pldn$^{f05716}$ flies bear a UAS construct in the pallidin gene allowing overexpression[5]. The elav-GeneSwitch stock was obtained from B Mollereau lab (Ecole Normal Supérieure de Lyon, France)[6]. For the induction by elav-GeneSwitch, Mifepristone was diluted in the food at a concentration of 50 µg/µL. For the induction with the TARGET system[7], flies were raised at 18°C, to allow inhibition of the Gal4-UAS induction system. Adult flies were then recorded at 18°C to assess sleep without induction, before being transferred to 30°C to trigger Gal4 activity. All lines were outcrossed 3 times to a Canton S reference strain or to other lines previously outcrossed to Canton S before being used for experimentation. The Gal4 and UAS parental lines outcrossed to Canton S served as genetic background controls in the experiments.

Sleep recording. Freshly hatched female flies were collected under $CO_2$ anesthesia and loaded individually at age 2-5 days into 5x65mm long glass tubes containing standard food medium. Sleep was recorded in a Light: Dark 12h:12h (LD) cycle at 25°C, 60% humidity using the Trikinetics DAMS system (www.trikinetics.com). Sleep parameters were evaluated across 3-6 days of baseline as described previously using 5 minutes immobility as criteria[8,9]. Sleep experiments were repeated at least 2 times. Distribution and homogeneity as well as statistical group comparisons were evaluated using the Microsoft Excel plugin software Statel. Kruskal-Wallis followed by posthoc comparisons were used in the statistical analysis. Mean ± SEM is plotted and the p value shown is the highest obtained among post hoc comparisons.
